# R allele of ACTN3 R577X gene polymorphism is associated with Elite Indian Boxer Status

**DOI:** 10.1101/2025.01.20.633833

**Authors:** Vijmendra Kumar Grover, Jai Prakash Verma, Ashish Kumar, Nivedita Sharma, Pramod Kumar Tiwari

## Abstract

Genetic variations are considered important for athletic performance. In this regard, polymorphisms in angiotensin I-converting enzyme (ACE) and α-actinin-3 genes are widely studied for their association with elite athlete status. The ACE gene regulates circulatory homeostasis, with the I variant of the ACE insertion/deletion (ACE I/D) gene polymorphism being associated with endurance performance, while the R allele of the R577X polymorphism of the ACTN3 gene, crucial for fast glycolytic muscle fibers, has been associated with speed and power performances. The present study investigated the association of these genetic variants with elite boxer (N=57) status and compared it with elite power/speed athletes (N=40), endurance athletes (N=44) and nonathletes (N=98) in the Indian population. The R allele frequency was found significantly higher in boxers than the nonathletes (*p* < 0.05). The allele and genotype frequencies of boxers, endurance and power/speed athletes did not differ significantly (*p* > 0.05). After analyzing overall athletic cohort on the basis of their level of performance, the frequency of RR genotype (24.1%) and R allele (47.7%) in national level athletes was found significantly higher than in nonathletes (8.7% and 34.2%, respectively), with the significant difference (*p* < 0.05) between national level endurance athletes and nonathletes only. The ACE I/D gene polymorphism was not found associated with any of the athletic cohorts (*p* > 0.05). Taken altogether, our study showed that the R allele of ACTN3 gene polymorphism is associated with elite boxer status as compared to the nonathletes.

**Highlights:**

- Boxers had a higher ACTN3 gene R allele frequency than nonathletes; genotype did not differ.
- Boxers with RR genotype had more than 3x higher odds versus nonathletes (*p* = 0.02) in the X dominant model (RR vs RX+XX).
- ACTN3 gene allele and genotype frequencies differed between overall athletic cohort and nonathletes.
- ACE gene frequencies showed no difference or association across athletic cohorts.

## 1. Introduction

Both genetic and non-genetic components contribute to the performance of an individual in any sport, with ∼66% of the variance in athlete status explained by additive genetic factors and the remaining due to nonshared environment (De Moor et al., 2007). Many studies have been conducted to identify the relationship of genetic factors with athletic performance (Semenova et al., 2023). The angiotensin I-converting enzyme insertion/deletion (ACE I/D) gene polymorphism and α-actinin-3 R577X (ACTN3) gene polymorphisms are the most studied genetic polymorphisms across different populations (Semenova et al., 2023). The α-actinin-3 is a structural protein of sarcomeric Z-line restricted to only fast glycolytic skeletal muscle fibers. C→T transition at position 1747 in exon 16 of this gene, converting arginine to a stop codon at residue 577 (R577X), results in the α-actinin-3 deficiency. This polymorphism in the ACTN3 gene influences normal variation in muscle function (Yang et al., 2003). On the other hand, the ACE gene is crucial for circulatory homeostasis in the renin-angiotensin system (RAS). It catalyzes the conversion of angiotensin I to vasoconstrictor angiotensin II and degradation of vasodilator bradykinin (Rigat et al., 1990). Insertion/deletion (I/D) of a 287 bp Alu sequence in intron 16 of the ACE gene is associated with circulating and tissue ACE levels, with higher ACE activity associated with the D variant, and lower ACE activity linked to the I variant. (Alvarez et al., 2000).

Earlier studies have shown that the I allele of the ACE gene and X allele of the ACTN3 gene are associated with endurance performance, whereas the D allele of the ACE gene and the R allele of the ACTN3 gene are more frequent in speed and power athletes (Ahmetov and Fedotovskaya, 2015). Most of these studies investigated associations with either endurance sport or power sport and only a limited number of studies analyzed the role of selected polymorphisms in sports of mixed nature (Eynon et al., 2014; Massidda et al., 2015; Semenova et al., 2023; Youn et al., 2021).

Boxing, one of the oldest and most popular combat sports, became part of the Summer Olympic program in 1904. Boxers need to maintain a high frequency of attacking movements to win in a boxing bout, even the defensive movements often end with a counterattack (Davis et al., 2013). Although aerobic power is essential to accelerate the recovery process within a boxing match, anaerobic power is linked with performance success in boxing (Chaabène et al., 2015). Boxers need to be quick and agile, able to move quickly around the ring, feint, and deliver fast and accurate strikes. Boxers need to be able to generate significant force with their punches in order to score points and knock out their opponents (Ruddock et al., 2018). Also, muscular power, a highly heritable characteristic, is essential for boxing performance (Chaabène et al., 2015; Zempo et al., 2017). Various studies have been conducted to understand the association of these athletic phenotypes with the ACTN3 R577X and ACE I/D gene polymorphisms (Semenova et al., 2023). However, no study has been conducted to investigate the association of the ACTN3 R577X and ACE I/D gene polymorphisms emphasizing boxers. Only a small number of boxers have been included as participants in combat sports group (Cocci et al., 2019; Franchini, 2014; Jung et al., 2016; Youn et al., 2021). Here, we have studied the ACTN3 R577X and ACE I/D gene polymorphisms in elite Indian boxers and compared it with the elite power and endurance athletes and nonathletic controls. In this study, we evaluated whether genetic variations in ACTN3 and ACE genes may be associated with the status of being an elite boxer.

## 2. Materials and methods

### 2.1 Participants

All the participants in the study were Indian elite athletes (boxers, endurance and power athletes) and nonathletes. Summary of participant demographics such as number and their age has been included in Supplementary Table 1. Data is presented as mean ± standard deviation (SD) for age. A total of 239 male participants (57 elite boxers, 44 elite endurance athletes, 40 elite power/speed athletes and 98 nonathletes) were selected for the study.

#### Criteria for elite athlete status

Those who had either won medals at senior national championship or had represented the national team at international tournaments of respective sport were defined as the elite level athletes. The selected athletes were categorized according to their best performance level into international level athletes (who have either represented India or won medals at world class tournaments like Asian championship, commonwealth championship, world championship, Asian games, commonwealth games, Olympic Games etc. in their respective sport) and national level athletes (other than the international level athletes).

#### Nonathletes

Nonathletes were those healthy individuals who did not have any competitive sport experience.

#### Categorization of sports

Endurance athletes belonged to events like 5,000 m and 10,000 m races, walking races, cross country and marathon races; power/speed athletes were those who participated in weight lifting, throwing events (shot put, discus throw, javelin throw), sprint races (100 m, 200 m, 400 m), hurdle races (110 m hurdles, 400 m hurdles), jumping events (long jump, high jump).

### 2.2 Sample collection and genotyping

Saliva samples were collected after obtaining a written consent as per the guidelines of the Institutional Human Research Ethics Committee, Jiwaji University, Gwalior.

For genetic analysis, DNA was isolated from the participants’ saliva samples containing buccal cells using the spin column based method, as per the manufacturer’s protocol (Qiagen blood mini kit). The integrity of DNA was checked onto 0.8% agarose gel electrophoresis (Figure 1). The ACTN3 and ACE gene sequences were amplified by polymerase chain reaction (PCR) using the specific primers (Supplementary Table 2). The amplified product of the ACE gene provided 477 bp fragment for the I allele and 190 bp fragment for the D allele. The reaction products were electrophoresed on 2% agarose gel and observed under the gel documentation system (Figure 2 (i)). For the determination of R577X polymorphism of the ACTN3 gene, PCR products were digested by DdeI restriction endonuclease for restriction fragment length polymorphism (RFLP). The digested products were separated on 3% agarose gel and observed under the gel documentation system. The DNA bands were of 205 bp and 86 bp for the R allele and 108 bp, 97 bp and 86 bp for X allele (Figure 2 (ii)).

**Figure 1.**
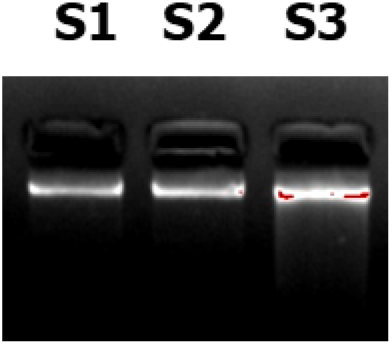
Integrity of gDNA. Integrity of gDNA under UV light onto 0.8% agarose gel in representative samples (S1- S3)

**Figure 2.**
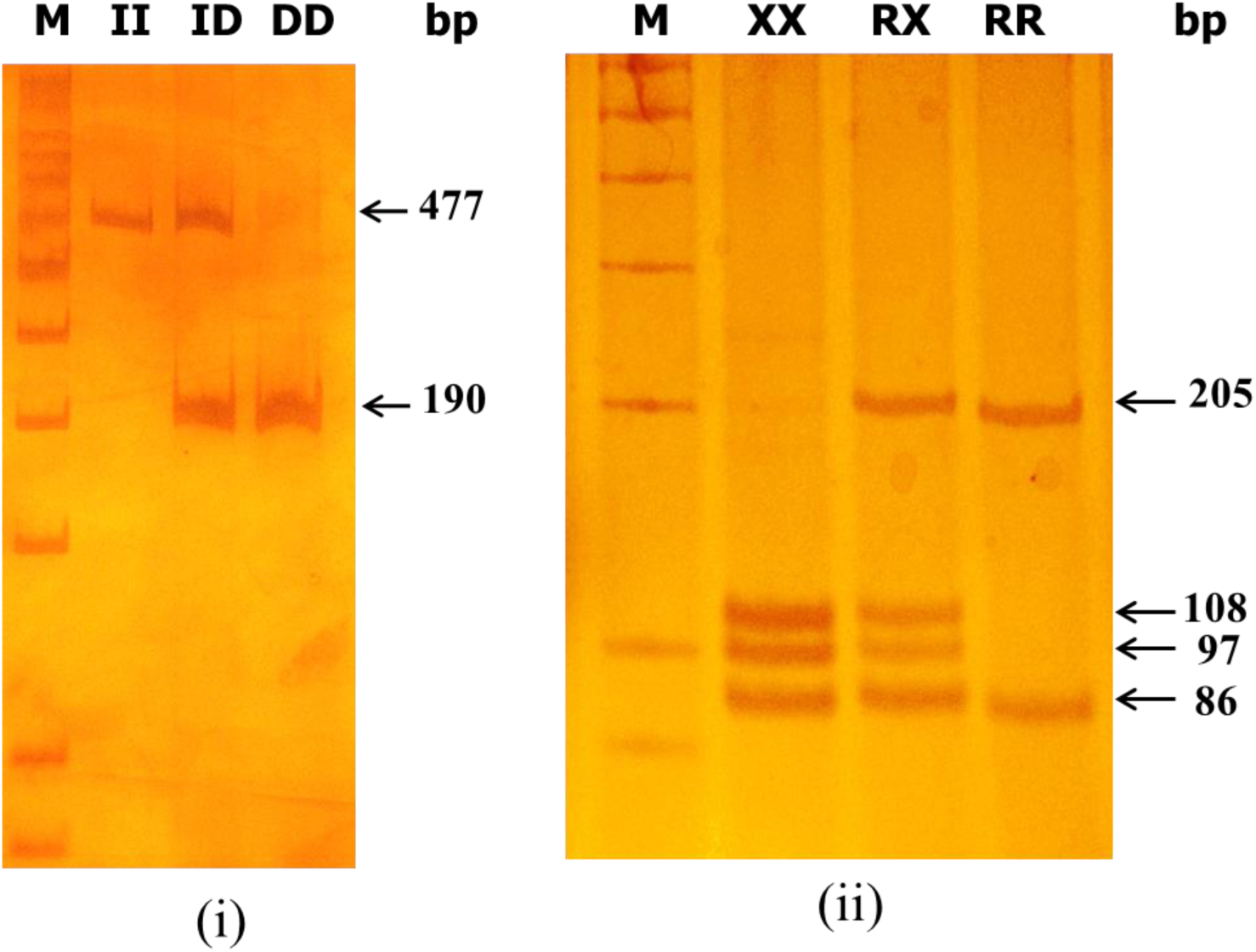
Images of ACE and ACTN3 gene polymorphisms. Agarose gel showing (i) genotypes for ACE I/D gene polymorphism (ii) genotypes for ACTN3 R577X gene polymorphism. Molecular size of ACE gene I allele is 477 bp and D allele is 190 bp. For the ACTN3 gene, the size of the R allele has 205 and 86 bp bands, and the X allele has 108, 97 and 86 bp bands after restriction enzyme DdeI digestion.

### 2.3 Statistical analyses

All the statistical analyses were done using SPSS 20.0 software. Allele frequencies were determined by gene counting. Genotype and allele frequencies between groups of athletes and controls were then compared by using the *χ2* test. Odds ratios (ORs) were analysed by logistic regression for athletic groups and nonathletes. The level of significance for all the statistical tests was set at 0.05.

## 3. Results

### 3.1 Distribution of ACTN3 R577X gene polymorphism

The genotypic data for ACTN3 R577X gene polymorphism showed significant differences in allele (*χ2* = 5.989; *p* = 0.016) and genotype frequencies (*χ2* = 7.302; *p* = 0.026) between overall athletic cohort and nonathletic controls as shown in Table 1 and 2, with the higher frequencies of R allele (45.7%) and RR genotype (21.6%) in athletes than that of nonathletes (34.2% and 8.7%, respectively) (Figure 3 (i) and (ii)). When the athletes were divided into boxers, sprint/power athlete and endurance athlete groups and compared with the nonathletes, the frequency of R allele in boxers (45.6%) was significantly higher (*χ2* = 3.843; *p* =0.050) than that in the nonathletes (34.2%). However, the genotype frequency of the boxers did not differ significantly as compared to the nonathletes (*χ2* = 5.899; *p* = 0.052). The allele and genotype frequencies of endurance and power/speed athletes did not differ significantly from nonathletes as shown in Table 2.

**Figure 3.**
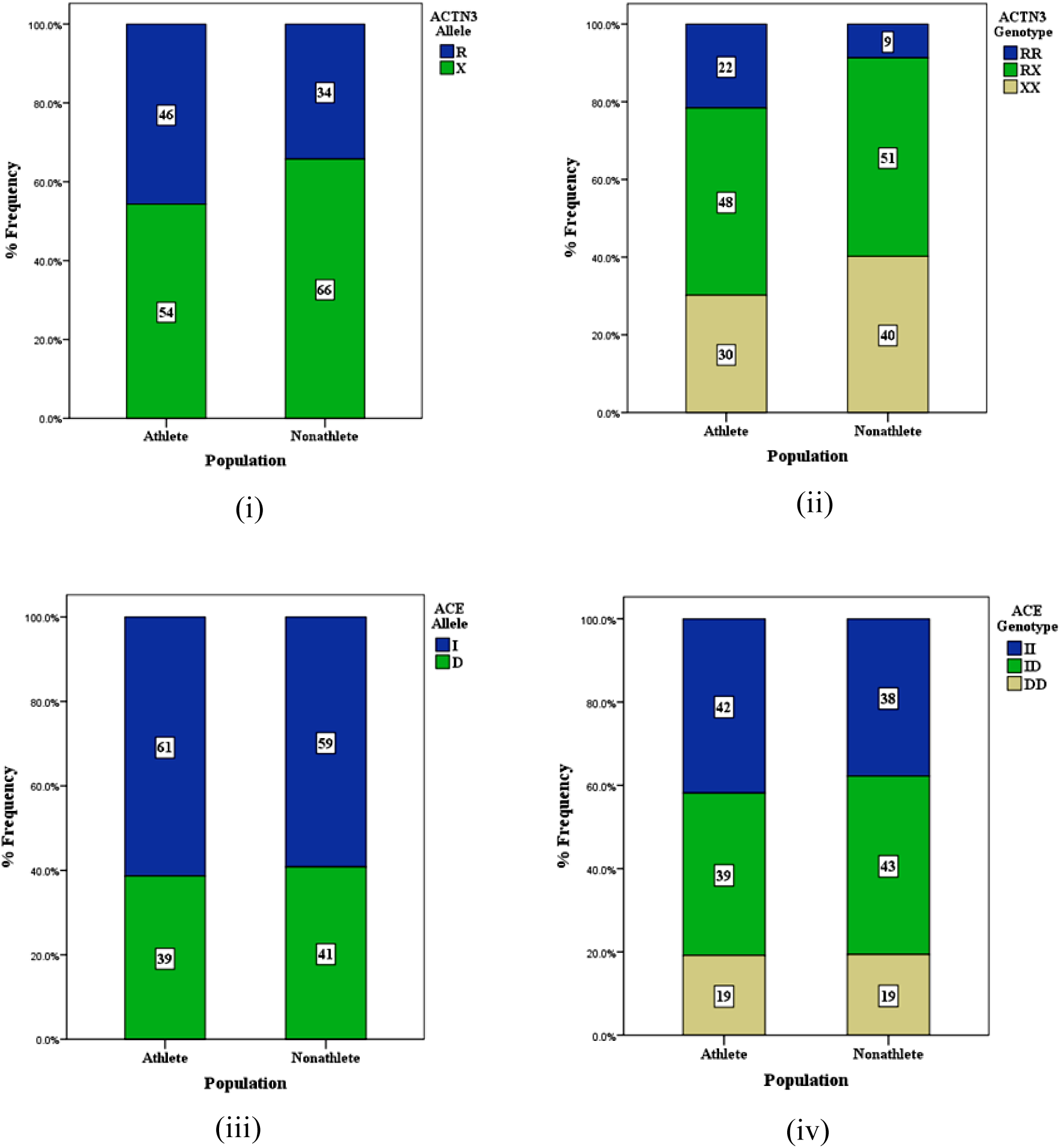
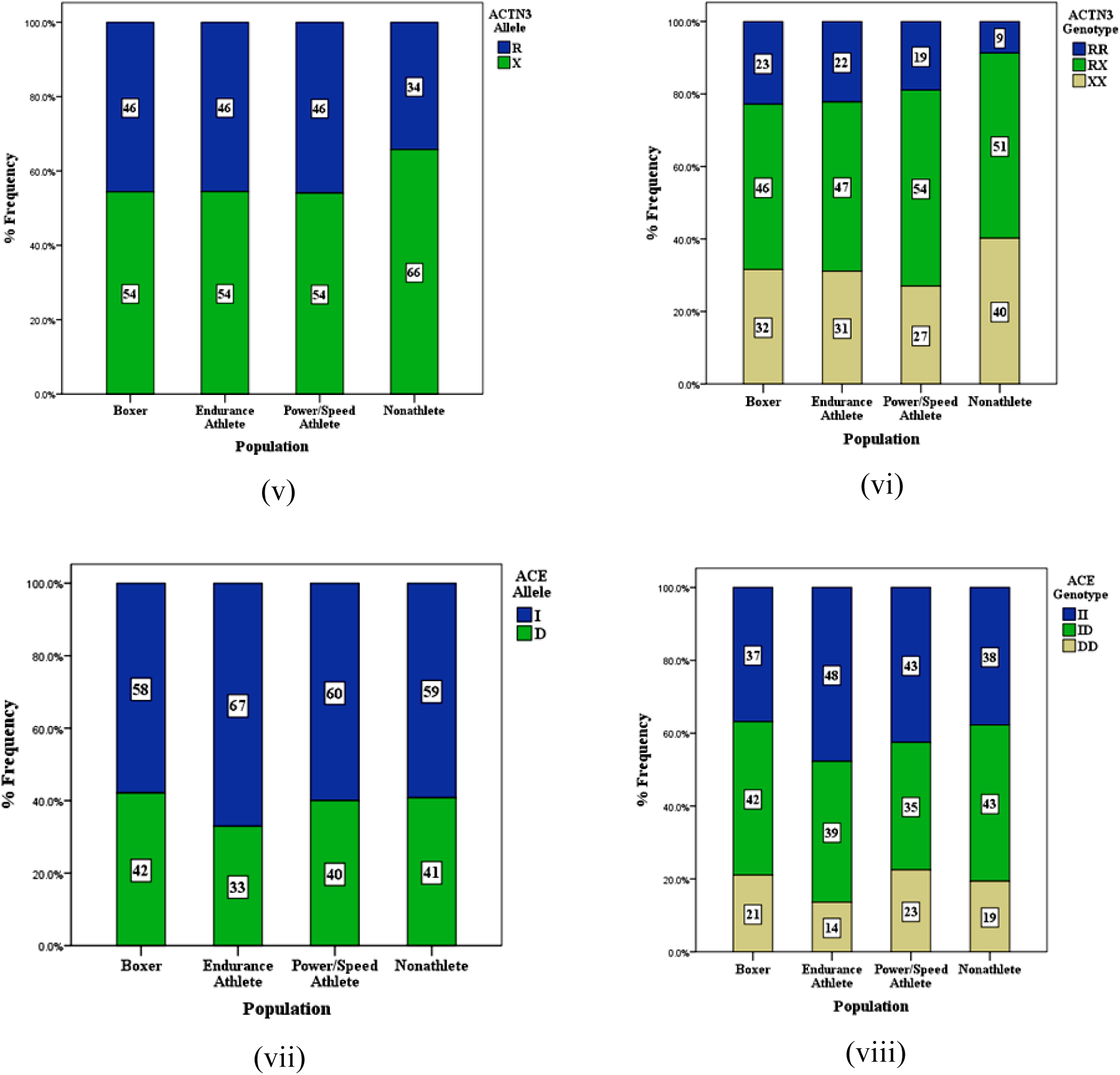
Graphs for distribution of ACE and ACTN3 gene polymorphisms. (i) Allele distribution of ACTN3 R577X gene polymorphism in overall athletic cohort and nonathletes. (ii) Genotype distribution of ACTN3 R577X gene polymorphism in overall athletic cohort and nonathletes. (iii) Allele distribution of ACE I/D gene polymorphism in overall athletic cohort and nonathletes. (iv) Genotype distribution of ACE I/D gene polymorphism in overall athletic cohort and nonathletes. (v) Allele distribution of ACTN3 R577X gene polymorphism in athletic groups and nonathletes. (vi) Genotype distribution of ACTN3 R577X gene polymorphism in athletic groups and nonathletes. (vii) Allele distribution of ACE I/D gene polymorphism in athletic groups and nonathletes. (viii) Genotype distribution of ACE I/D gene polymorphism in athletic groups and nonathletes.

**Table 1.** Genotype and allele distribution of ACE I/D and ACTN3 R577X gene polymorphisms.

|  | **ACTN3 Genotype** |  |  | **ACTN3 Allele** |  |
| --- | --- | --- | --- | --- | --- |
| **Group** | **RR** | **RX** | **XX** | **R** | **X** |
| Overall Athletic Cohort | 30 (21.6%) | 67 (48.2%) | 42 (30.2%) | 127 (45.7%) | 151 (54.3%) |
| Boxer | 13 (22.8%) | 26 (45.6%) | 18 (31.6%) | 52 (45.6%) | 62 (54.4%) |
| Endurance | 10 (22.2%) | 21 (46.7%) | 14 (31.1%) | 41 (45.6%) | 49 (54.4%) |
| Power/Speed | 7 (18.9%) | 20 (54.1%) | 10 (27.0%) | 34 (45.9%) | 40 (54.1%) |
| Nonathletes | 8 (8.7%) | 47 (51.1%) | 37 (40.2%) | 63 (34.2%) | 121 (65.8%) |
|  | **ACE Genotype** |  |  | **ACE Allele** |  |
| **Group** | **II** | **ID** | **DD** | **I** | **D** |
| Overall Athletic Cohort | 59 (41.8%) | 55 (39.0%) | 27 (19.1%) | 173 (61.3%) | 109 (38.7%) |
| Boxer | 21 (36.8%) | 24 (42.1%) | 12 (21.1%) | 66 (57.9%) | 48 (42.1%) |
| Endurance | 21 (47.7%) | 17 (38.6%) | 6 (13.6%) | 59 (67.0%) | 29 (33.0%) |
| Power/Speed | 17 (42.5%) | 14 (35.0%) | 9 (22.5%) | 48 (60.0%) | 32 (40.0%) |
| Nonathletes | 37 (37.8%) | 42 (42.9%) | 19 (19.4%) | 116 (59.2%) | 80 (40.8%) |

**Table 2.** Chi-square analysis of ACTN3 gene genotypes and alleles across athletic groups and nonathletes.

| Comparison Groups | Category | Genotype<br>($\chi^2$ value) | Genotype<br>(*p*-value) | Allele<br>($\chi^2$ value) | Allele<br>(*p*-value) |
| --- | --- | --- | --- | --- | --- |
| **Athletes vs. Nonathletes** | All Athletes | 7.302 | 0.026 | 5.989 | 0.016 |
|  | Boxers | 5.899 | 0.052 | 3.843 | 0.050 |
|  | Endurance | 5.0 | 0.082 | 3.287 | 0.070 |
|  | Power/Speed | 3.677 | 0.159 | 3.083 | 0.079 |
| **National vs. International Athletes** | National vs International | 0.928 | 0.629 | 0.763 | 0.382 |
|  | National vs. Nonathletes | 8.426 | 0.015 | 6.711 | 0.010 |
|  | International vs. Nonathletes | 2.597 | 0.273 | 1.853 | 0.173 |
| **Athletic Groups by National vs. International** | Boxers - National vs. International | 0.216 | 0.898 | 0.044 | 0.834 |
|  | Boxers - National vs. Nonathletes | 4.981 | 0.083 | 3.418 | 0.064 |
|  | Boxers - International vs. Nonathletes | 3.240 | 0.198 | 1.219 | 0.270 |
|  | Endurance - National vs. International | 1.050 | 0.592 | 0.560 | 0.454 |
|  | Endurance - National vs. Nonathletes | 6.474 | 0.039 | 3.827 | 0.050 |
|  | Endurance - International vs. Nonathletes | 0.465 | 0.792 | 0.376 | 0.540 |
|  | Power/Speed - National vs. International | 0.503 | 0.778 | 0.416 | 0.519 |
|  | Power/Speed - National vs. Nonathletes | 3.940 | 0.139 | 3.069 | 0.080 |
|  | Power/Speed - International vs. Nonathletes | 1.160 | 0.560 | 0.977 | 0.323 |

When the athletes were divided into international and national level groups based on their performance level, the genotype (*χ2* = 8.426; *p* = 0.015) and allele frequency (*χ2* = 6.711; *p* = 0.010) differed significantly between all national level athletes and nonathletes, with genotype frequency of RR genotype, 24.1% and allele frequency of R allele, 47.7% in national level athletes and 8.7% and 34.2%, respectively in nonathletes (Table 2). Furthermore, only the national level endurance athletes differed significantly from nonathletes in genotype distribution (*χ2* = 6.474; *p* = 0.039) and allele distribution (*χ2* = 3.827; *p* = 0.050), shown in Table 2.

### 3.2 Distribution of ACE I/D gene polymorphism

The genotypic data showed that the boxers and power/speed athletes had a higher proportion of the DD genotype (21.1% and 22.5%, respectively) than the nonathletes (19.4%), though the difference was not significant. The endurance and power/speed athletes had a higher proportion of the II genotype (47.7% and 42.5%, respectively) than the nonathletes (37.8%) but the difference was not significant (shown in Table 3).

**Table 3.** Chi-Square analysis of ACE gene genotypes and alleles across athletic groups and nonathletes.

| Comparison Groups | Category | Genotype ($\chi^2$ value) | Genotype ($p$-value) | Allele ($\chi^2$ Value) | Allele ($p$-value) |
| --- | --- | --- | --- | --- | --- |
| **Athletes vs. Nonathletes** | All Athletes | 0.454 | 0.797 | 0.226 | 0.636 |
|  | Boxers | 0.063 | 0.969 | 0.049 | 0.824 |
|  | Endurance | 1.440 | 0.487 | 1.587 | 0.208 |
|  | Power/Speed | 0.731 | 0.694 | 0.016 | 0.900 |
| **National vs. International Athletes** | National vs International | 1.441 | 0.487 | 1.425 | 0.233 |
|  | National vs. Nonathletes | 0.063 | 0.969 | 0.012 | 0.913 |
|  | International vs. Nonathletes | 1.558 | 0.459 | 1.265 | 0.261 |
| **Athletic Groups by National vs. International** | Boxers - National vs. International | 0.262 | 0.877 | 0.298 | 0.585 |
|  | Boxers - National vs. Nonathletes | 0.195 | 0.907 | 0.201 | 0.654 |
|  | Boxers - International vs. Nonathletes | 0.077 | 0.962 | 0.080 | 0.777 |
|  | Endurance - National vs. International | 2.384 | 0.304 | 1.440 | 0.230 |
|  | Endurance - National vs. Nonathletes | 0.388 | 0.824 | 0.199 | 0.655 |
|  | Endurance - International vs. Nonathletes | 3.484 | 0.175 | 2.904 | 0.088 |
|  | Power/Speed - National vs. International | 0.357 | 0.837 | 0.134 | 0.715 |
|  | Power/Speed - National vs. Nonathletes | 0.944 | 0.624 | 0.022 | 0.882 |
|  | Power/Speed - International vs. Nonathletes | 0.211 | 0.900 | 0.106 | 0.744 |

### 3.3 The odds ratio of ACTN3 R577X genotypes and ACE I/D genotype

The results of logistic regression are summarized in table 4 in the form of ORs. The combined athletic group had 2.89 (*p* = 0.012) times higher odds of harbouring the RR genotype than nonathletes in the X dominant model (RR vs RX+XX). The OR for a boxer harbouring the RR genotype in X dominant model (RR vs RX+XX) compared with nonathlete was 3.10 (*p* = 0.02). The endurance athletes had the OR of 3.00 (*p* = 0.03) for harbouring the RR genotype in X dominant model (RR vs RX+XX) as compared to nonathletes. The ORs were higher for an athlete (irrespective of athletic category), boxer, endurance athlete and power/speed athlete with RR or RX genotype in the R dominant model (RR+RX vs XX) as compared to nonathlete. However, the ORs in the I dominant model (II+ID vs DD) and D dominant model (II vs ID+DD) did not suggest any relationship for any of the cohort.

**Table 4.** Odds ratios (ORs) of genotypes for athletes and control participants according to sport type.

| Group | Athlete vs Nonathlete |  | Boxer vs Nonathlete |  | Endurance vs Nonathlete |  | Power vs Nonathlete |  |
| --- | --- | --- | --- | --- | --- | --- | --- | --- |
| | OR (95% CI) | $p$-value | OR (95% CI) | $p$-value | OR (95% CI) | $p$-value | OR (95% CI) | $p$-value |
| RR+RX vs XX | 1.554<br>(0.894- 2.701) | 0.118 | 1.458<br>(0.726- 2.926) | 0.289 | 1.490<br>(0.699- 3.173) | 0.302 | 1.817<br>(0.787- 4.194) | 0.162 |
| RR vs RX+XX | 2.889<br>(1.259- 6.633) | 0.012 | 3.102<br>(1.196- 8.048) | 0.020 | 3.001<br>(1.093- 8.236) | 0.033 | 2.450<br>(0.818- 7.336) | 0.109 |
| II+ID vs DD | 1.015<br>(0.529- 1.949) | 0.963 | 0.902<br>(0.470- 1.733) | 0.803 | 1.524<br>(0.563- 4.124) | 0.408 | 0.829<br>(0.338- 2.028) | 0.680 |
| II vs ID+DD | 1.186<br>(0.700- 2.010) | 0.526 | 0.962<br>(0.489- 1.891) | 0.910 | 1.505<br>(0.734- 3.089) | 0.265 | 1.219<br>(0.577- 2.575) | 0.605 |

## 4. Discussion

In the present study, we investigated the allelic and genotypic frequencies of ACTN3 R577X and ACE I/D gene polymorphisms in elite boxers and compared them with elite endurance athletes, power/speed athletes and nonathletes. We observed a higher frequency of R allele in the boxers. The odds of a boxer harbouring the RR genotype (RR vs RX+XX) are more than thrice as compared to the nonathlete. However, the genotypic and allelic frequencies of ACE gene I/D polymorphism was not different from that of the nonathletes. Moreover, the genotypic profile for both the genes of the boxers was not different from that of the Indian endurance athletes and power/speed athletes.

Regarding ACE I/D gene polymorphism, II genotype or I allele of ACE I/D gene polymorphism has been associated with endurance athlete status (Collins et al., 2004; Gayagay et al., 1998; Lucía et al., 2005; Myerson et al., 1999) and D allele or DD genotype with power or sprint performance (Nazarov et al., 2001; Papadimitriou et al., 2008; Scott et al., 2010; Wang et al., 2013). However, the association of I allele with endurance athlete status was not significant when it was analyzed in Kenyans and East African runners (Ash et al., 2011; Scott et al., 2005). The D allele and DD genotype of ACE gene were also not significant in Jamaican sprinters (Scott et al., 2010). These populations rule the world in their respective athletic events, but these genes did not show prevalence there. Negative or no associations with ACE genotypes are reported in other populations as well (Ahmetov and Fedotovskaya, 2015). In the present study, the ACE gene did not show any association across any of the athletic cohort, suggesting that ACE I/D gene polymorphism is not a genetic factor that differs in elite Indian athletes.

Boxing is a mixed power/endurance sport, relying on a combination of bioenergetic pathways (Barley et al., 2019; Chaabène et al., 2015). The nature of this sport differentiates it from the athletic activities used in this study for comparison, which are either predominantly anaerobic or aerobic. Given the high physiological demands of elite boxing, studying gene polymorphisms of ACE and ACTN3 genes might provide deeper understanding of the association between performance and genetics. However, none of the available studies specifically examined elite boxers, making direct comparisons difficult (Anastasiou et al., 2024; Franchini, 2014; Youn et al., 2021). Therefore, for comparison, we used results from other studies involving other combat sports, as their physiological demands match most closely with boxing.

When compared between Japanese judo athletes and control, no association was found for the genotype and allele frequencies of the ACE I/D polymorphism and judo status or endurance (Itaka et al., 2016). In elite Italian male athletes, no statistically significant differences in genotype distribution or allele frequencies for ACE polymorphism were found when comparing combat sport athletes, motorcycle riders and soccer players (Cocci et al., 2019). However, the ACE genotype distribution and allele frequency of Japanese wrestlers significantly differed from those of the controls (Kikuchi et al., 2012).

To get success in boxing, boxers need to throw repetitive high intensity punches which requires well-developed muscle strength, muscle power, anaerobic power and capacity (Cocci et al., 2019). In skeletal muscles, α-actinin-3 protein (expressed in humans with RR or RX genotypes) is restricted to fast glycolytic fibres and constitute the predominant protein component of the sarcomeric Z line, where it stabilises the muscle contractile apparatus, providing a mechanical advantage in strength and power performances over ACTN3 deficient (XX) humans (MacArthur and North, 2004). In addition to their mechanical role, ACTN3 might play an important role in the determination of fiber type distribution. A positive association was found between the amount, surface area, percentage of fast glycolytic fibres (IIx) and ACTN3 R allele in humans (Vincent et al., 2007). The hypothesis that the R allele enhances high velocity muscle tasks was supported by the study conducted by Vincent et al. (2007).

The higher allelic frequency of the R allele in Indian boxers than that in the nonathletes aligns with findings observed in other combat athletic populations. The frequency of the ACTN3 R allele was significantly higher in Japanese wrestlers than in controls (*p* < 0.05) (Kikuchi et al., 2013). In an investigation by Akazawa et al. (2022), the frequency of RR genotype was also higher in martial arts athletes (*p* < 0.05) compared to control. Although, in elite Italian male athletes, no statistically significant differences in genotype distribution or allele frequencies of ACTN3 polymorphism were found when comparing combat sport athletes with other athletic populations (Cocci et al., 2019). Similarly, the present study also showed insignificant differences in the genotypes of ACTN3 polymorphism in boxers, endurance athletes and power/speed athletes.

Analysis of the other athletic subgroups in the present study showed that Indian endurance athletes differed significantly from nonathletes for allele and genotype frequencies, with higher frequency of R allele and RR genotype in endurance athletes than nonathletes. The hypothesis supporting the advantage of ACTN3 deficiency in endurance performance has also not been supported in other athletic populations (Ahmetov et al., 2010; Lucia et al., 2006; Niemi and Majamaa, 2005; Paparini et al., 2007; Saunders et al., 2007; Yang et al., 2007)Ahmetov et al., 2008; Ahmetov et al., 2010; Lucia et al., 2006; Niemi and Majamaa, 2005; Paparini et al., 2007; Saunders et al., 2007; Yang et al., 2007). Moreover, the ACTN3 R577X gene polymorphism was not significantly associated with the status of being an elite power/speed athlete in our study. Similar negative associations of RR genotype or R allele with sprint/power athlete status was found in Lithuanian, Italian and Nigerian cohorts (Ginevičienė et al., 2011; Sessa et al., 2011; Yang et al., 2007).

## 5. Conclusion and recommendations

Taken together, boxers were found to have the higher frequency of R allele of ACTN3 R577X gene polymorphism and national level endurance athletes had higher frequencies of R allele and RR genotypes than nonathletes. ACE I/D gene polymorphism is not a significant genetic factor in Indian elite athletes.

The present study has genotyped two of the key candidate genes for athletic performance in a cohort of Indian elite athletes. Research on the relevance of these genetic variants for performance phenotype has contradictory findings across different populations (Semenova et al., 2023). This might be due to the dependence of performance on multiple factors and its polygenic nature or because of the epigenetic influences. In addition to the two genetic variations studied, other variants have been identified as potentially significant, and require further investigations (Semenova et al., 2023; Youn et al., 2021). Some studies investigated the associations between gene polymorphisms and traits such as personality, social behavior, mood, arousal, pain tolerance and others in combat sports athletes (Anastasiou et al., 2024; Youn et al., 2021). Future research should also consider genetic markers associated with other sport-related phenotypes, utilizing approaches such as GWAS, whole-genome sequencing, epigenetic, transcriptomic, proteomic and metabolomic profiling and meta-analyses in large cohorts of athletes, before these findings can be extended to practice in sport (Trent and Yu, 2009). In recent years, there has been a growing market for direct-to-consumer (DTC) tests claiming to identify athletic talents in children primarily based on these two gene polymorphisms, owing to the large number of studies conducted on these genes (Webborn et al., 2015). However, the current understanding about genetics of athletic performance is limited, highlighting the need for further investigations to assess the role of genetic polymorphisms in identifying sports talent. Future research should focus on establishing more precise relationships between genetic profiles and athletic abilities. Such research has the potential to improve the talent identification and development process by providing a deeper understanding of the genetic factors that influence performance.

## CRediT authorship contribution statement

**Vijmendra Kumar Grover:** Conceptualization, Methodology, Sample Collection, Analysis, Writing – original draft. **Jai Prakash Verma:** Conceptualization, Supervision, Resources, Writing - review & editing. **Ashish Kumar:** Methodology, Sample Collection, Writing – original draft. **Nivedita Sharma:** Methodology, Analysis. **Pramod Kumar Tiwari:** Conceptualization, Supervision, Resources, Writing - review & editing.

## Acknowledgement

We sincerely thank all the study participants for donating sample and time to facilitate this study. The invaluable assistance provided by the Molecular and Human Genetics department in conducting all experiments is acknowledged. This work was supported by the scholarship provided by University Grants Commission (UGC), India. VKG was the recipient of Senior Research Fellowship and AK was recipient of Dr. D.S. Kothari Postdoctoral Fellowship from UGC.

## Supplementary Material

**Supplementary Table 1.** Demographic characteristics of study participants.

| Group | Number | Age in years (M ± S.D.) |
| --- | --- | --- |
| **Overall Athletic Cohort** | 141 | 28.5 ± 5.95 |
| **International Level** | 54 | 29.5 ± 5.87 |
| **National Level** | 87 | 27.8 ± 5.93 |
| **Boxers** | 57 | 29.6 ± 7.19 |
| **International Level** | 17 | 28.4 ± 6.73 |
| **National Level** | 40 | 32 ± 7.62 |
| **Endurance Athletes** | 44 | 29.3 ± 4.47 |
| **International Level** | 16 | 28.6 ± 5.02 |
| **National Level** | 28 | 30.4 ± 3.15 |
| **Power/Speed Athletes** | 40 | 25.6 ± 4.07 |
| **International Level** | 21 | 25 ± 4.47 |
| **National Level** | 19 | 26.1 ± 3.69 |
| **Nonathletes** | 98 | 26.1 ± 4.30 |

**Supplementary Table 2.**
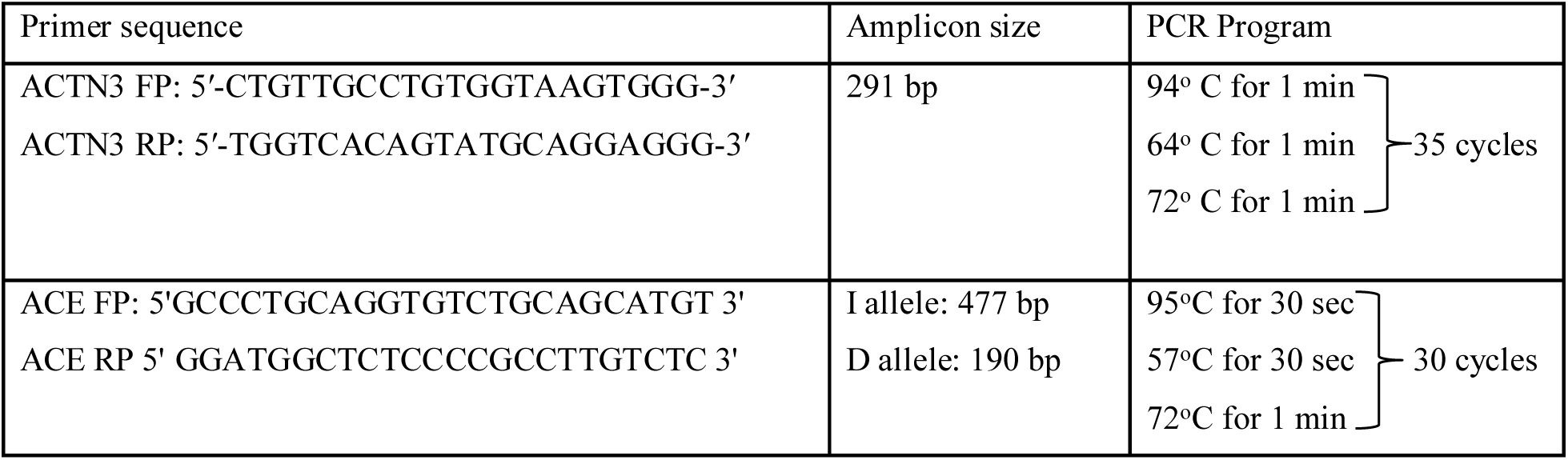
Details of the primers and PCR program used for the study.

